# The pseudokinase Trib1 regulates the transition of exhausted T cells to a KLR^+^ CD8^+^ effector state and its deletion improves checkpoint blockade

**DOI:** 10.1101/2023.02.16.528833

**Authors:** Susan E. McClory, Oishi Bardhan, Kelly S. Rome, Josephine R. Giles, Amy E. Baxter, Lanwei Xu, Phyllis A. Gimotty, Robert B. Faryabi, E. John Wherry, Warren S. Pear, Martha S. Jordan

**Affiliations:** Division of Oncology, Children’s Hospital of Philadelphia, Philadelphia, PA, USA; Department of Pathology and Laboratory Medicine, Abramson Family Cancer Research Institute, Perelman School of Medicine, University of Pennsylvania, Philadelphia, PA, USA; Department of Pathology, Children’s Hospital of Philadelphia, Philadelphia, PA, USA; Department of Systems Pharmacology and Translational Therapeutics, University of Pennsylvania, Philadelphia, PA, USA; Institute for Immunology, Perelman School of Medicine, University of Pennsylvania, Philadelphia, PA, USA; Parker Institute for Cancer Immunotherapy, Perelman School of Medicine, University of Pennsylvania, Philadelphia, PA 19104, USA; Department of Biostatistics, Epidemiology and Informatics, Perelman School of Medicine, University of Pennsylvania, Philadelphia, PA, USA

## Abstract

T cell exhaustion (T_EX_) impairs the ability of T cells to clear chronic infection or cancer. While exhausted T cells are hypofunctional, some exhausted T cells retain effector gene signatures, a feature that is associated with expression of KLRs (killer lectin-like receptors). Although KLR^+^ T cells may improve control of chronic antigen, the signaling molecules regulating this population are poorly understood. Using scRNA-seq, flow cytometry, RNA velocity, and scTCR-seq, we demonstrate that deleting the pseudokinase Trib1 shifts T_EX_ towards CX3CR1^+^ intermediates (T_INT_) with robust enrichment of KLR^+^CD8^+^ T cells (T_KLR_) via clonal T cell expansion. These changes are associated with globally increased KLR gene expression throughout the exhaustion program. Further, Trib1 loss augments anti-PD-L1 blockade to improve viral clearance by expanding the T_KLR_ population. Together, these data identify Trib1 as an important regulator of T cell exhaustion whose targeting enhances the KLR^+^ effector state and improves the response to checkpoint inhibitor therapy.

## Introduction

T cell exhaustion is a homeostatic state that limits the immune system’s ability to eradicate chronic antigen such as cancer or infection. During the initial stages of chronic antigen presence, T cells have a robust effector response. Eventually proliferation and function dampen, and T cells settle into a state of exhaustion, marked by increased expression of inhibitory receptors, decreased cytokine production, and poor expansion ^1-3^. However, T cells retain some effector function, and in the best-case scenario, T cell exhaustion serves to limit immune pathology. Yet, exhaustion can be problematic in the setting of cancer or chronic infection because antigen is never cleared, and disease progresses ^1-3^. Thus, there is a strong interest in developing T cell-based therapies that target T cell exhaustion to overcome the dampened immune response in chronic disease.

In CD8^+^ T cells, exhaustion occurs through a stepwise developmental process whereby a naïve T cell encounters chronic antigen and differentiates into an exhausted cell precursor ^4^ and then progenitor (T_PROG_), which ultimately produces terminally exhausted T cells (T_TERM_). The progenitor cells express high levels of Ly108 and TCF1 ^2,4-6^, they can self-renew, and proliferate in response to PD1 blockade ^7-11^. T_PROG_ give rise to transitional or intermediate cells (T_INT_) that express CX3CR1 and are the cells that accumulate following PD1 inhibition. These cells retain T cell function and are important for viral and tumor control ^4,6,10,12,13^. T_INT_ further differentiate into terminally exhausted cells (T_TERM_), expressing high levels of inhibitory receptors including PD1, Tim3, TIGIT, and Lag3, as well as CD69, CD39, and CD101 receptors. T_TERM_ do not self-renew or expand, have decreased function and are relatively short-lived ^2,3,6,10,13^. In addition to T_TERM_, recent evidence supports that T_INT_ can also differentiate into a T_KLR_ population ^4,14,15^. T_KLR_ are a potentially important population of effector-like T cells, which express high levels of KLRG1 in addition to Killer Lectin-like Receptors (KLRs), such as NKG2D (encoded by *Klrk1*), NKG2A (encoded by *Klrc1*), CD94 (encoded by *Klrd1*), NK1.1 (encoded by *Klrb1c*) and other proteins expressed by both effector T cells and NK cells, including granzymes, the transcription factors Zeb2, Tbx21 (T-bet) and ID2, and the chemokine receptor CX3CR1 with low CXCR6 expression ^4,14-16^. In the setting of chronic infection, T_KLR_ have transcriptional profiles similar to effector T cells from acute viral infections but express higher amounts of PD1 and TOX (a TF known to enforce the T_EX_ state) than do effector CD8^+^ T cells during acute infection ^4,14^. The findings that T_INT_ can differentiate into either T_KLR_ or T_TERM_ cells ^4,14,15^, suggest that T_KLR_ may be an alternative to terminally exhausted cells and expansion of this population may improve outcomes during chronic antigen exposure. Indeed, effector-like CD8^+^ T cells expressing KLRs, such as NK1.1, NKG2D, NKG2A, and CD94 have been described in both mouse and human models of disease, including chronic infection, cancer, and autoimmunity ^17-20^. They are potent effector cells with strong cytotoxic potential and may play a role in self-recognition of stressed, infected, transformed, or hyperactive immune cells. This raises the question of whether the fate of CD8^+^ T cells during chronic antigen exposure can be shifted away from a terminal T_EX_ and towards a more effector-like T_KLR_ to improve anti-viral or anti-tumor immunity and immune therapies.

We previously identified the pseudokinase Tribbles 1 (Trib1) as a novel regulator of T cell effector function and activation during chronic infection^16^. Trib1 is a member of the Tribbles pseudokinase family (Trib) that is present in multicellular organisms^21^. Tribs are defined by a central pseudokinase domain that is important for protein interactions and a C-terminal domain that interacts with the E3 ligase Cop1 and is critical for regulating Trib-mediated protein degradation. Important targets of Trib1-mediated protein degradation in mammalian cells are C/EBP family members^22,23^. We previously identified a MALT1 binding site in the N-terminus of Trib1 that appears to be unique to Trib1 among Trib proteins^16,24^. The TRIB1:MALT1 interaction prevents MALT1 from forming the CBM (CARMA1/BCL10/MALT1) complex, which is important for transducing TCR signals^16^. In the setting of chronic LCMV (clone 13) infection, Trib1 mRNA is increased in both CD4^+^ and CD8^+^ T cells. Using CD4^cre^Trib1^f/f^ mice (Trib1 KO) and CD4^cre^Trib1^+/+^ mice (WT), we demonstrated that Trib1 KO mice infected with LCMV clone 13 had more KLRG1^+^CD8^+^ effector-like cells, their CD8^+^ and CD4^+^ T cells were more robust cytokine producers, and Trib1 KO mice had lower viral titers after 30 days of infection compared to WT mice^16^. Recently, Trib1 was found to be highly expressed during T cell exhaustion in healthy human donor peripheral blood and has multiple accessible chromatin regions (ACRs) in exhausted human tumor infiltrating lymphocytes (TILS)^25^, suggesting that Trib1 not only regulates T cell exhaustion in mouse LCMV, but is a highly relevant gene in human T cell exhaustion as well. Thus, we sought out to better understand how the pathway of T cell exhaustion can be shifted towards a more functional CD8^+^ T cell fate by a molecular regulator, such as Trib1, and to better understand if targeting Trib1 could improve immunotherapy for cancer or infectious disease. Here, we demonstrate that Trib1 deletion from T cells during chronic infection increases expression of KLR and KLR-associated genes throughout the exhausted CD8^+^ T cell subsets and promotes clonal expansion of T_INT_ and T_KLR_ subsets. Additionally, Trib1 KO improves the response to immune checkpoint blockade (ICB) by expanding the T_KLR_ subset in contrast to PD-L1-blockade alone, where this subset is not expanded. Thus, inhibiting Trib1 has the potential to expand the T_INT_ and T_KLR_ populations, and in doing so, increase the benefit of ICB.

## Results

### Trib1 restrains KLR gene expression throughout the T cell exhaustion pathway

Previous scRNA sequencing of all CD44^+^TCR*β*^+^ splenocytes from WT and Trib1 KO mice infected with LCMV clone 13 for 15 days revealed that a CD8^+^KLRG1^+^ population was preferentially enriched in Trib1 KO cells compared to WT T cells, and that this population had a transcriptional profile similar to effector T cells from acute infection ^16^. To gain more granularity within the CD8^+^ T population, we reanalyzed this data set excluding cells expressing *Cd4* mRNA. The remaining *Cd4*-low cells expressed high *Cd8* mRNA with no *Cd4* mRNA, whereas the excluded cells expressed high *Cd4* and low *Cd8* mRNA demonstrating effective isolation of CD8^+^ T cells from this data set (**Fig. S1A**). To better understand how KLR associated genes are regulated by Trib1 throughout T cell exhaustion, we used TooManyCells ^26^ to generate a CD8^+^-specific dendrogram with 9 branches or clusters (**Fig. 1A**). This tree expressed uniform *Cd3e, Cd8a* and *Cd8b* mRNA (**Fig. S1B**). We identified nine clusters and used differential gene expression to annotate the tree (**Figs. 1A, S1B-C**). Cluster 1 expressed high levels of *Klrg1* and *Cx3cr1*; in addition, there was prominent expression of multiple additional KLRs, such as Clusters 6 and 7 expressed *Klrb1c, Klrc1, Klrd1, Klre1*, and *Klrk1*. The widespread expression of multiple KLRs suggests that this population represents a newly termed “T_KLR_” subset during chronic infection ^4,14^. This cluster also expressed transcripts characteristic of effector cells such as granzymes and the transcription factors *Id2* and *Zeb2*. Clusters 2 and 3 expressed *Cx3cr1* but low levels of KLRs, consistent with T_INT_. The primary difference between clusters 2 and 3 was higher expression of proliferation genes by cluster 3 (**Fig. S1D**). When these proliferation genes were regressed and cell identities projected onto the other clusters, cluster 3 cells most closely resembled cluster 2, confirming they were proliferating CX3CR1^+^ T_INT_ (**Fig. S1E**). Cluster 4 expressed high levels of terminal exhaustion genes (*Tox, Pdcd1, Lag3, Tigit)*, consistent with a T_TERM_ phenotype (**Fig. S1B-C**). Clusters 6 and 7 expressed *Tcf7* and *Slamf6*, but low levels of *Tox, Pdcd1, Lag3*, and *Tigit* consistent with early T_PROG_ cells. Cluster 8 expressed *Tcf7* and *Slamf6*, but also higher levels of *Tox, Pdcd1, Lag3*, and *Tigit*, also consistent with T_PROG_ cells. Cluster 5 was a small population comprising only 138 cells (out of 5,113 total) and had features that overlapped with progenitors (expression of *Slamf6* and *Tcf7*), but also had features shared with multiple populations (**Fig. S1B-C**). Cluster 9 had a high proliferative gene signature and when proliferation genes were removed from the analysis and the resultant DEGs were regressed onto the tree, cluster 9 had a signature most closely resembling clusters 7, 8 and 3, suggesting these proliferative cells largely were composed of cells transitioning from a T_PROG_ to a T_INT_ identity (**Fig. S1F**). Of note, GSEA comparing our data to the LCMV-specific populations identified by Giles et al ^4^ at day 15 of clone 13 infection revealed strikingly similar patterns (**Fig. S1G**). Together, these data reinforce the identify of known T_EX_ subsets: T_PROG_, T_INT_ and T_TERM_ and highlight the presence of an effector-like T_KLR_ subset characterized by high expression of effector-associated and KLR genes.

**Figure 1:**
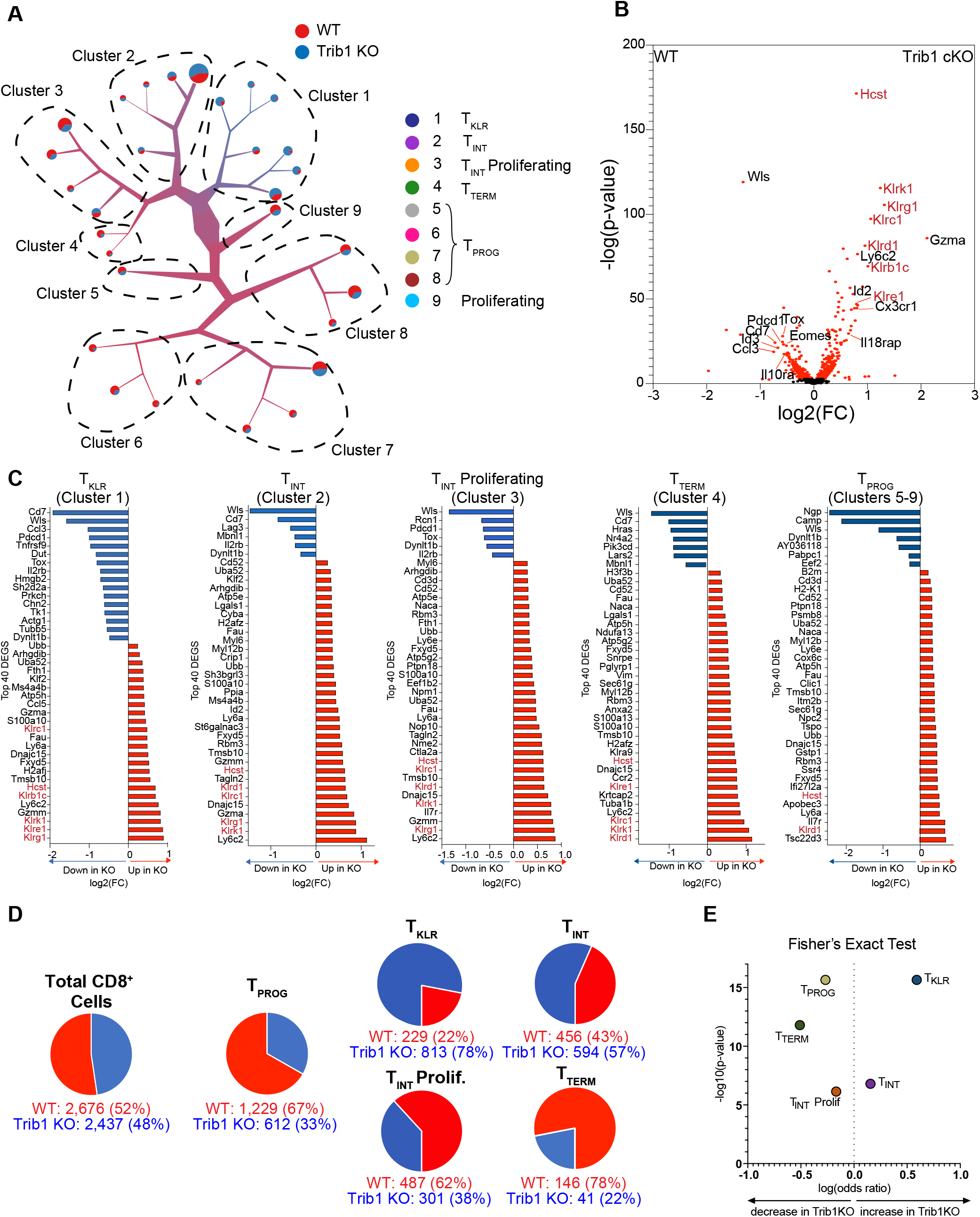
Trib1 deletion during chronic LCMV infection is characterized by enrichment of KLR-associated genes within a complete pathway of T cell exhaustion. scRNA-sequencing was performed on FACS-sorted total activated T cells (CD44^+^TCR*β*^+^) from Trib1 WT (CD4-Cre^+^Trib1^+/+^) or Trib1 KO (CD4-Cre^+^Trib1^f/f^) mice that had been infected with clone 13 15 days prior, as previously described^16^. To exclude CD4^+^ T cells from this new analysis, all barcodes expressing any *CD4* mRNA were excluded. (A) A new visualization of single-cell clustering of all CD8^+^ cells using TooManyCells ^26^. With this method, individual cells all begin at the central node and then are recursively divided based on transcriptional differences, resulting in branches or clusters on the tree. Branch width corresponds to cell number and color represents the proportion of cellular genotype making up that branch. In this case, blue represents Trib1 KO and red Trib1 WT. Each terminal branch ends with a node or leaf that is sized based on the number of cells present in that node. 9 main branches or clusters are identified as shown in (A). (B) Volcano plot depicting DEGs in CD8^+^ cells of either Trib1 WT or Trib1 KO cells. The KLRs and the *Hcst* (encodes DAP10 a critical adaptor protein for NKG2D signaling) are highlighted in red text. (C) The top 40 DEGs between Trib1 WT and Trib1 KO CD8^+^ cells in Cluster 1-4 and as combined progenitor populations (Clusters 5-9). Blue indicates higher expression in WT CD8^+^ cells and red higher expression in Trib1 KO CD8^+^ cells. (D) Proportion of WT and Trib1 KO cells in the representative populations, as defined by the scRNA-seq using TooManyCells. T_PROG_ cell frequency was calculated by combining cell numbers from clusters 5, 6, 7, and 8 on the dendogram in (A). T_KLR_, T_INT_, T_INT Proliferating_, and T_TERM_ correspond to clusters 1, 2, 3, and 4 respectively. (B) Fisher’s exact test demonstrating differential representation of WT and Trib1 KO cells in T_PROG_, T_KLR_, T_INT_, T_INT Proliferating_, and T_TERM_ subsets. Trib1 WT: CD4-cre^+^ Trib1^+/+^, Trib1 KO: CD4-cre^+^ Trib1^F/F^

Using this CD8^+^ tree, we identified differentially expressed genes (DEGs) between Trib1 KO and WT cells. Trib1 KO CD8^+^ cells were specifically enriched for multiple KLRs such as *Klrg1, Klrk1*(encoding NKG2D), *Klrc1* (encoding NKG2A), *Klrd1* (encoding CD94), *Klrb1c* (encoding NK1.1), *Klre1* (encoding NKG2I), and KLR-associated genes such as *Hcst* (encoding DAP10, a signaling adaptor for NKG2D). They also expressed higher levels of *Gzma* and *Cx3xr1* than WT cells, and lower levels of exhaustion-associated genes such as *Pcdc1, Tox*, and *Eomes* (**Fig. 1B**) suggesting that absence of Trib1 biases towards an activated phenotype. Furthermore, when Trib1 KO cells were compared to WT cells within each cluster, KO cells consistently had higher KLR expression at all stages of the exhaustion program (**Fig. 1C)**, suggesting that Trib1 regulates KLR gene expression during chronic infection.

We next looked at the distribution of Trib1 KO and WT cells in each CD8+ T_EX_ subset. The overall contribution of WT and Trib1 KO cells to the analysis was similar (52% and 48%, respectively) (**Fig. 1D**). T_PROG_ (clusters 5-8) were enriched for WT cells (67%) compared to Trib1 KO cells (33%), whereas T_INT_ (clusters 2 and 3) was relatively evenly distributed between the two genotypes. In contrast, T_TERM_ (cluster 4) was strongly enriched for WT cells (78%) compared to Trib1 KO cells (22%) and conversely, the T_KLR_ subset (cluster 1) was enriched for Trib1 KO cells which accounted for 78% of this cluster compared to 22% contributed by WT cells **(Fig. 1D-E)**. These data indicate that deleting Trib1 pushes T_PROG_ towards T_INT_ and shifts the population of exhausted CD8^+^ T cells away from T_TERM_ and towards the more effector-like T_KLR_ population.

### Trib1-deficient CD8^+^ T cells are enriched for CX3CR1^+^ cells largely due to a significant expansion of CX3CR1^+^KLRG1^+^ T_KLR_ cells

We next investigated whether the effects of Trib1 deletion on the exhaustion pathway were observed at the cellular level in LCMV-specific CD8^+^ cells. WT and Trib1 KO mice were infected with clone 13 for 27-30 days, and splenocytes were analyzed for LCMV-specific CD8^+^ cells using a Gp33 tetramer. CX3CR1 and Ly108 expression during chronic infection can be used to identify CD8^+^ T_INT_ cells and T_PROG_ cells, respectively ^4,10,13,14^. Thus, we analyzed the expression of these proteins on Trib1 WT and KO cells. Total Gp33^+^CD8^+^CX3CR1^+^ cells were expanded in Trib1 KO compared to WT mice, accompanied by a relative decrease in Gp33^+^CD8^+^Ly108^+^ T_PROG_ cells (**Figs. 2A, S2A-B)**. Similar to the scRNA-seq data (**Fig. 1**), total CX3CR1^+^ cells can be divided based on KLRG1 expression: CX3CR1+ KLRG1^−^ cells were consistent with cluster 2 and 3 T_INT_ cells that expressed high levels of *Cx3cr1* but low levels of *Klrg1*; CX3CR1^+^KLRG1^+^ cells corresponded to the cluster 1 T_KLR_ subset (**Figs. 2B, S2C-D**). Loss of Trib1 resulted in both a relative and absolute increase in LCMV-specific CX3CR1^+^KLRG1^+^ T_KLR_ cells, compared to WT mice (**Figs. 2B, S2D**). At this timepoint, Trib1 KO mice had more Gp33^+^ T_INT_ cells, defined by CX3CR1^+^KLRG1^−^ expression (although the magnitude of increase did not always reach statistical significance) (**Figs. 2B, S2C**). This shift towards both CX3CR1^+^ populations was accompanied by a proportional decrease in CD101^+^ T_TERM_ cells in Trib1 KO mice compared to WT (**Figs. 2C, S1E**). Additionally, the Gp33^+^CD8^+^KLRG1^+^ cells identified at this timepoint also co-express CD94 and NKG2A, further confirming T_KLR_ features (**Fig. 2D**). These data confirm the scRNA-seq data at the protein level, suggesting that Trib1 regulates the terminal fate of CD8^+^ T cell exhaustion by restraining differentiation into T_KLR_ and that deletion of Trib1 results in a robust skewing towards a more effector-like T_KLR_ phenotype. Furthermore, these data reveal that loss of Trib1 enhances the T_INT_ pool, a population known to be critical for viral control.

**Figure 2:**
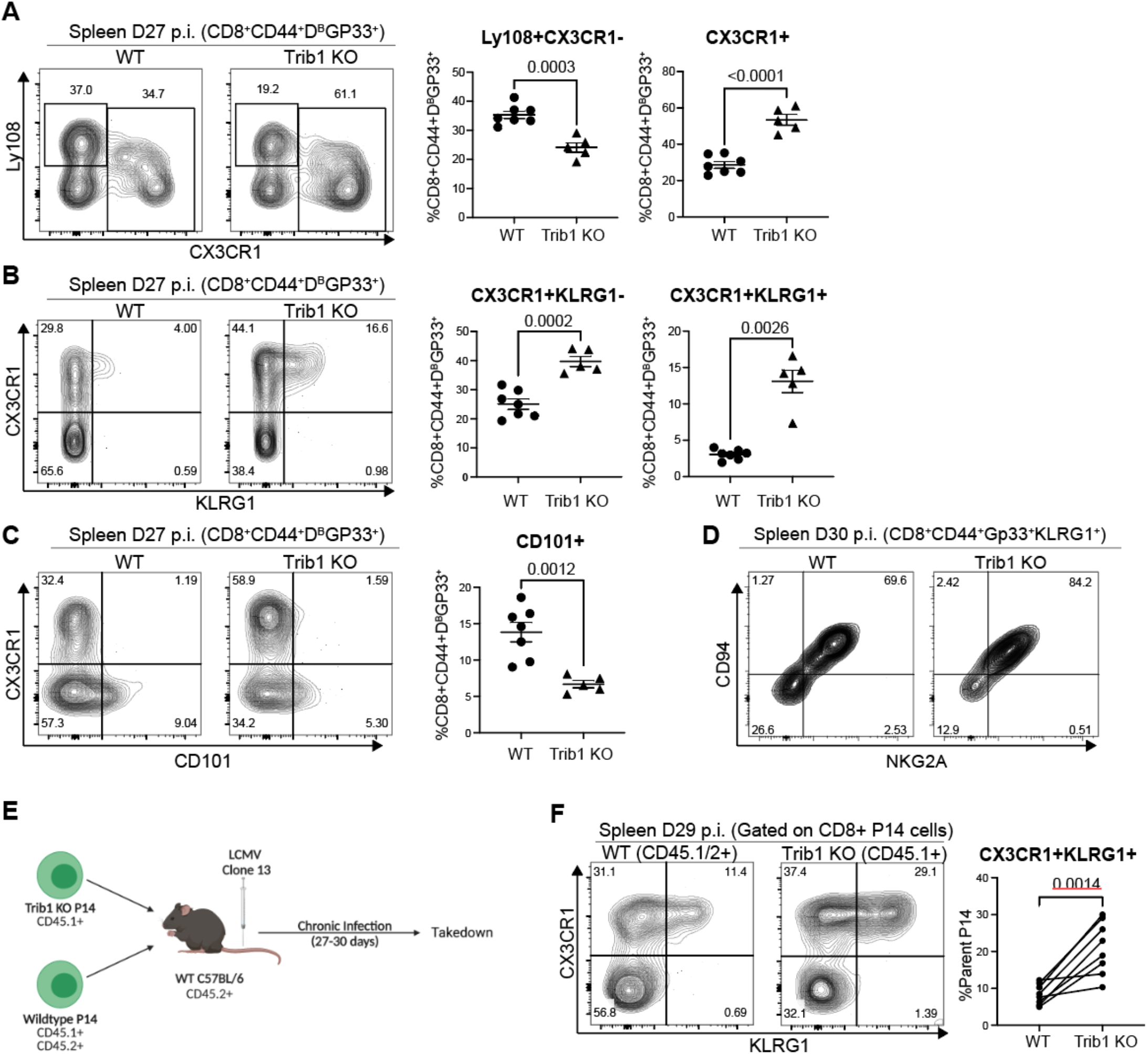
Trib1 deletion drives CD8^+^ T cells towards a KLR^+^ effector-like fate during chronic infection. (A) Representative flow plots with the proportion of Ly108^+^ T_PROG_ and Total CX3CR1^+^ cells in WT or Trib1 KO mice infected with LCMV clone 13 for 27-30 days. Cells are gated on CD8^+^CD44^+^Gp33-Tetramer^+^ splenocytes. (B) Representative flow plots of CD8^+^CD44^+^Gp33-Tetramer^+^ cells that are either CX3CR1^+^KLRG1^-^ T_INT_ or CX3CR1^+^KLRG1^+^ T_KLR_ at day 27 in the spleen. (C) Representative flow plots with the proportion of CD8^+^CD44^+^Gp33-Tetramer^+^ cells that are CD101^+^ T_INT_ at day 27 in the spleen. (D) Representative CD94 and NKG2A expression on CD8^+^CD44^+^Gp33-Tetramer^+^KLRG1^+^ T_KLR_ cells. (E) Experimental design for P14 adoptive transfer experiments. (F) Representative flow plots of WT P14 cells (gated on CD45.1/2+ events) or Trib1 KO P14 cells (gated on CD45.1+ events) that are either CX3CR1^+^KLRG1^-^ T_INT_ or CX3CR1^+^KLRG1^+^ T_KLR_ at day 29 in the spleen. Data in (A-C) are representative of 6-7 experiments with 5-8 mice per genotype. Data in (D) are representative of 1 experiment with 5-7 mice. Data in (F) are representative of 2 experiments with 8-10 recipient mice per experiment. Error bars are mean ± SEM. P values for (A-D) are calculated using either Student’s t test or unpaired t test with Welch’s correction based on variance between genotypes. P values for (F) are calculated using a paired T test given that both KO and WT P14 cells were transferred into the same individual mice. Trib1 WT: CD4-cre^+^ Trib1^+/+^, Trib1 cKO: CD4-cre^+^ Trib1^F/F^

To determine if the skewing of CD8^+^ T cell fate towards the T_KLR_ subset in Trib1 KO mice was CD8-intrinsic, we co-transferred WT and Trib1 KO LCMV-specific TCR transgenic T cells (P14 cells) into congenically distinct recipient mice that were subsequently infected with clone 13 (**Figs. 2E-F**). This approach was important because loss of Trib1 in CD4^+^ T cells increased their proliferation and cytokine production ^16^, which could impact CD8^+^ T cell exhaustion through CD8-extrinsic mechanisms. Analysis of donor P14^+^ cells 29-30 days post-infection revealed a significant skewing to CX3CR1^+^KLRG1^+^ T_KLR_ Trib1 KO cells compared to WT P14 cells, indicating an intrinsic role for Trib1 in regulating the T_KLR_ state (**Fig. 2F**). This skewing was less robust but still significant when comparing absolute numbers **(Fig. S2F)** due to differences in overall recovery of WT versus Trib1 KO P14 cells (**Fig. S2G**). This finding suggests that Trib1 expression in CD4^+^ cells may play a role in the persistence of Trib1 KO CD8^+^ cells. Together, these data indicate that Trib1 deletion from CD8^+^ T cells specifically and intrinsically promotes a T_KLR_ fate.

### Trib1 deletion shifts the pathway of T cell exhaustion towards a T_KLR_ fate

To better determine how Trib1 influences the T cell differentiation pathway during chronic antigen exposure, we used RNA velocity to predict the directional flow of cell differentiation states within the CD8^+^ cells shown in **Fig. 1**. RNA velocity predicts the future fates of individual cells based on RNA content and splicing states ^27-29^. Cell identities of each cluster are shown in **Fig. 3A**. Cluster 9 was omitted from the analysis given its high proliferative gene signature and that its DEG profile overlaps significantly with other cell types when proliferative genes are removed (**Fig. S1D**). RNA velocity was calculated using Velocyto ^28^ and after visualization of CD8^+^ cells from either WT or Trib1 KO mice by uniform manifold approximation and projection (UMAP) using scanPY, RNA velocity was embedded onto both UMAPs by Cellrank ^29^ and scvelo ^28^ (**Fig. 3B-C**). Pruned partition-based graph abstraction (PAGA) was used to show the predicted connectivity and directional flow of clusters 1-8 (**Fig. 3D-E**). Using these visualizations, both WT and Trib1 KO CD8^+^ T cells had cell states that begin with Clusters 6 and/or 7 T_PROG_ and progressed towards Cluster 8 and to a smaller extent Cluster 5, in keeping with Clusters 6 and 7 being earlier T_PROG_. In both WT and Trib1 KO, T_PROG_ (Clusters 5, 6, 7, and 8) progressed towards Clusters 2 and 3 (T_INT_). WT cells then bifurcated towards either Cluster 4 (T_TERM_) or Cluster 1 (T_KLR_). While Trib1 KO CD8^+^ cells were also capable of differentiating into T_KLR_ or T_TERM_, the propensity to differentiate towards T_KLR_ was much stronger, as evidenced by both the directional flow predicted by RNA velocity as well as the visible frequency of T_KLR_ in the Trib1 KO UMAP (**Fig. 3D-E**). These data suggest that the T_INT_ stage can be a bifurcation point in the T_EX_ pathway with the potential to become either T_TERM_ or T_KLR_. Furthermore, our data demonstrate that T cell specific Trib1 deletion does not alter the path of T cell exhaustion; however, the absence of Trib1 promotes the trajectory of T_INT_ cells towards T_KLR_ cell state. Thus, these data support a model of CD8^+^ T cell exhaustion that begins with differentiation into a T_PROG_ and then T_INT,_ which can further differentiate into either T_KLR_ or T_TERM_, and that this branchpoint is regulated by Trib1 (**Fig. 3F-G)**.

**Figure 3:**
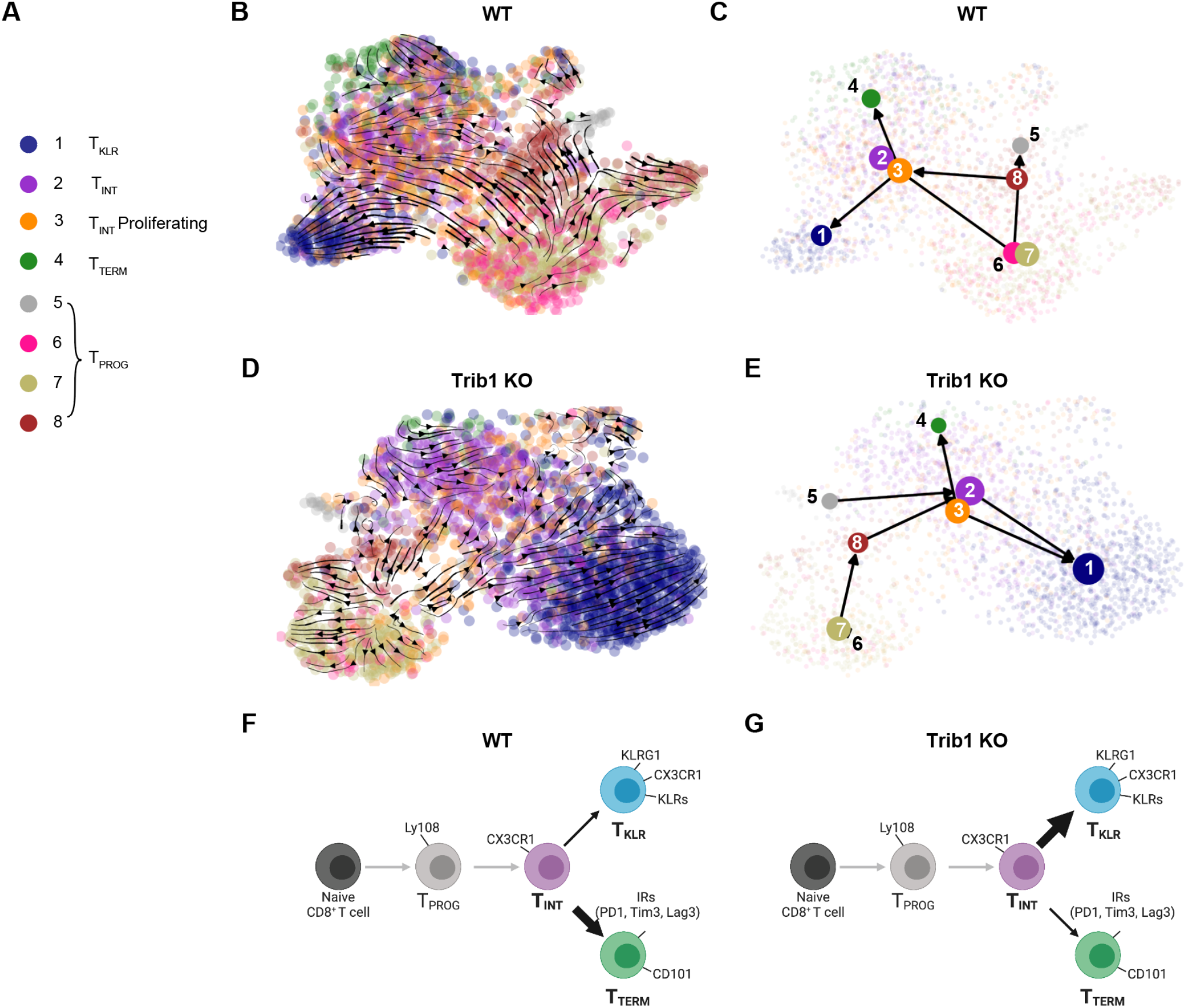
Trib1 KO shifts the pathway of T cell exhaustion towards a KLR^+^ fate as demonstrated by RNA Velocity. (A) Cell clusters identified in Figure 1 by TooManyCells. Cluster 9 has been excluded as it contains gene expression profiles of several other clusters, but with high expression of proliferative genes. (B) WT and (D) Trib1 KO RNA Velocity as projected on UMAP using the scRNA-seq data in Fig 2. Color of circles indicates the cluster identity. Arrows depict the directional flow of cell identities based on RNA Velocity as calculated by Velocyto and projected by Cellrank and scvelo. (B) Trib1 WT and (E) Trib1 KO pruned partition-based graph abstraction (PAGA) showing predicted connectivity of clusters 1-8. (F-G) Proposed model of T cell exhaustion in the WT (F) and Trib1 KO (G) setting. WT: CD4-cre^+^ Trib1^+/+^, Trib1 KO: CD4-cre^+^ Trib1^F/F^

### Accumulation of T_KLR_ cells in Trib1 KO mice is accompanied by clonal expansion during chronic infection

Recent reports suggest that T_TERM_ and T_KLR_ are terminal differentiation states that develop from a common CX3CR1^+^ T_INT_ cell during chronic infection ^4,14,15^, and that TCR clonal behaviors can trace the expansion and differentiation of single exhausted T cells into either of these two states ^14,15^. To investigate the clonal behavior and ontogeny of Trib1 KO CD8^+^ T cells during chronic infection, we performed paired single cell TCR sequencing with scRNA-seq of CD8^+^CD44^+^ T cells from WT and Trib1 KO mice 15 days after LCMV clone 13 infection. 2,904 TCR sequences (clonotypes) were identified in the CD8^+^ T cells of WT mice and 1,858 clonotypes in Trib1 KO mice. Of these, the majority were unexpanded single cell clones (91% in WT and 81% in Trib1 KO), reflecting a polyclonal CD8^+^ T cell pool. However, both WT and Trib1 KO mice had a small, but significant proportion of TCR clonotypes with more than 2 cells per sequence. Of these expanded clones, WT clonotypes ranged from 2-32 cells and Trib1 KO ranged from 2-138 cells (**Fig. 4A**). We determined the subset distribution of the 20-most expanded TCR clonotypes of each genotype. The majority of the 20-most expanded WT clonotypes fell into several cell subsets including T_PROG_, T_INT_, T_TERM_, and T_KLR_, whereas the top 20 Trib1 KO clonotypes were found more selectively in the T_KLR_ and T_INT_ subsets (**Fig. 4B**). Indeed, the top 20 clonotypes from Trib1 KO cells were most abundantly enriched for T_INT_ and T_KLR_, suggesting that Trib1 regulates the clonal expansion of these two populations presumably starting at the T_INT_ state given that the expanded clonotypes are first detected in abundance in the T_INT_ and then shared with the T_KLR._ (**Fig. 4B)**. From these data, clonotype behaviors can be described based on the cell subset distribution within a specific clonotype. Specifically, we asked if Trib1 deletion shifts the pathway of T_EX_ by impacting the clonal expansion of two alternative fates: T_KLR_ or T_TERM._ Using a 4-cell cutoff for significantly expanded clones, clonotype behaviors can be divided into 4 general categories: **Divergent clonotypes** contain both terminal subsets (T_KLR_ and T_TERM_), **T**_**KLR**_**-biased** contain T_KLR_ but not T_TERM_, **T**_**TERM**_**-biased** contain T_TERM_ but not T_KLR_, and **T**_**INT**_ contain neither T_KLR_ nor T_TERM_ (**Fig. 4C**). There were no significantly expanded clonotypes that contained only T_PROG_ cells in either WT or Trib1 KO cells. Notably, Trib1 KO cells had an over-representation of T_KLR_-biased clonotype behaviors, whereas WT cells had more T_TERM_-biased behaviors (**Fig. 4C**). Furthermore, Trib1 KO cells had a robust expansion of T_KLR_-biased clonotypes that contained only T_KLR_ and T_INT_ cells (**Fig 4C**). Expansion of Trib1 KO T_INT_ cells suggests Trib1 functions prior to the last step of transition from T_INT_ to T_KLR._ Together, these data indicate that in the absence of Trib1, there is clonal expansion of T cells shared between the T_INT_ and T_KLR_ subsets, suggesting that Trib1 specifically regulates the accumulation of these two subsets.

**Figure 4:**
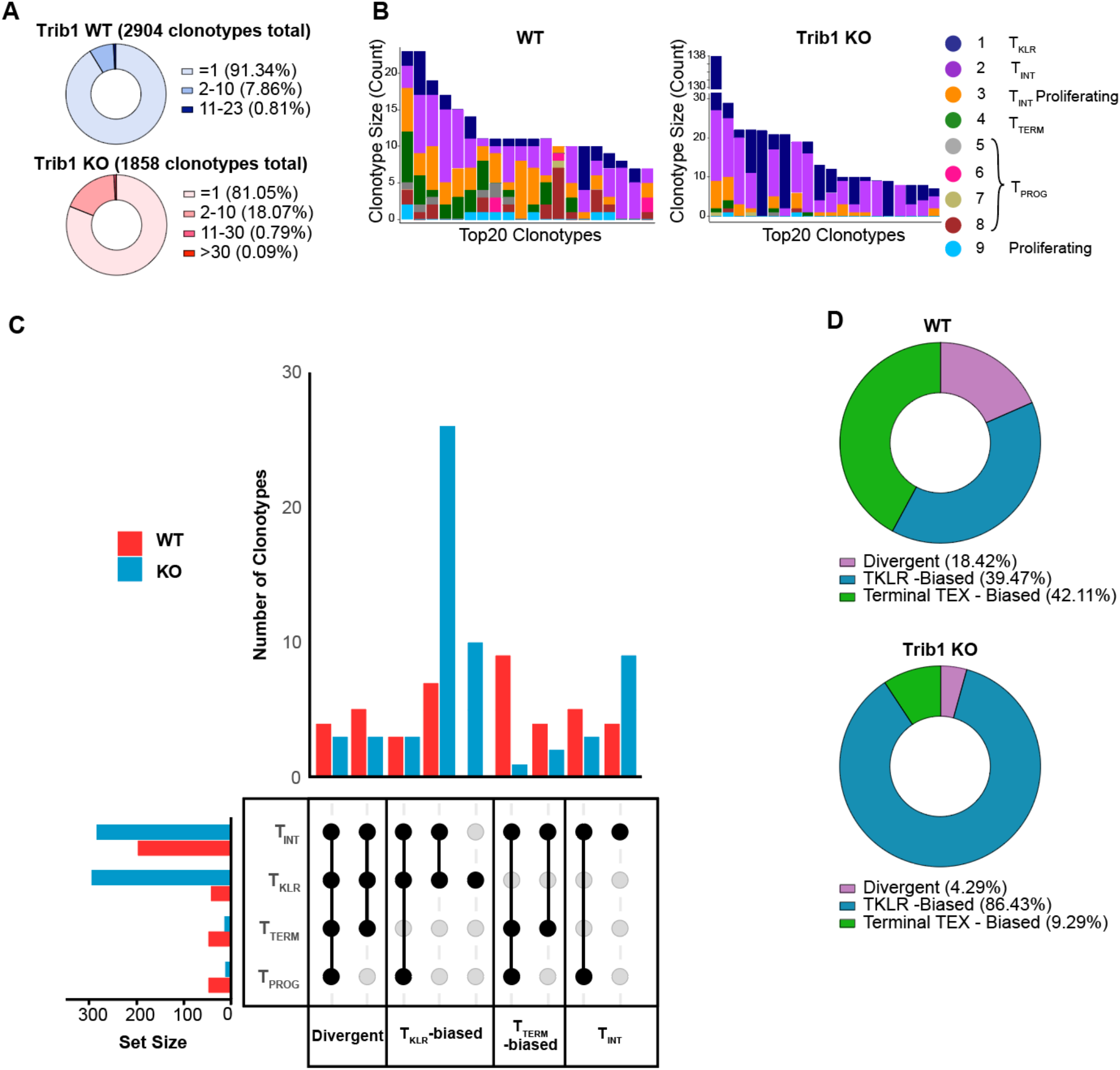
Trib1 KO drives clonal expansion of KLR^+^ T cells. (A) Total clonotypes from Trib1 WT and Trib1 KO cells containing 1 cells, 2-10 cells, 11-30 cells, or >30 cells. (B) Top 20 TCR sequences (clonotypes) identified in WT and Trib1 KO cells. Colors indicate presence of that clonotype in the corresponding cell cluster. (C) Upset plots depicting clonotype behavior (unique combinations of four generalized clusters) of WT (red) or Trib1 KO (blue) cells. Set size bar plots indicate the total number of cells in each cluster in all significantly expanded clones. d) Frequency of clonotypes that are T_KLR_-biased (blue), T_TERM_-biased (green), or Divergent (purple).

We next considered the breakdown of three alternative fates within the entire WT or Trib1 KO genotypes: T_KLR_-biased, T_TERM_-biased, or Divergent as defined above. WT mice contained a relatively equal percentage of T_KLR_-biased (39.47%) and terminal T_EX_-biased (42.11%) clonotypes, and a slightly smaller percentage of Divergent clonotypes (18.42%) (**Fig. 4D**). In contrast, Trib1 KO cells contained a much larger proportion of T_KLR_-biased clonotypes (86.43%) than either of the other two terminal fate behaviors (T_TERM_-biased: 9.29%; Divergent: 4.29%) (**Fig. 4D**). Taken together, these data show that Trib1 deletion from the T cell compartment shifts differentiation towards a T_KLR_ fate, and that this skewing is accompanied by clonal expansion of T_INT_ and T_KLR_ subsets.

### Trib1 deletion augments PD1 blockade

Efficacy of PD1 blockade depends on anti-PD1-mediated expansion of T_PROG_ cells, which then differentiate into T_INT_ and subsequently expand ^7-9,13^. We previously demonstrated that loss of Trib1 is sufficient to decrease viral load during chronic viral infection ^16^. Given the impact of Trib1 deletion on increasing effector-like differentiation and enhancing the T_INT_ pool, we hypothesized that disrupting Trib1 expression could further improve the efficacy of PD1 blockade. To test this possibility, WT and Trib1 KO mice were infected with clone 13 and treated with 5 doses of anti-PDL1 antibody or PBS between days 16 and 28 p.i. **(Fig. 5A**). Consistent with our previous report, Trib1 KO mice had lower viral titers than WT at day 29 p.i. (**Fig. 5B**). Importantly, the combination of Trib1 KO and PD1 blockade resulted in the lowest viral titers and were significantly lower than WT mice treated with PD1 blockade (**Fig. 5B**), demonstrating that deletion of Trib1 from T cells augments ICB to improve viral clearance.

**Figure 5:**
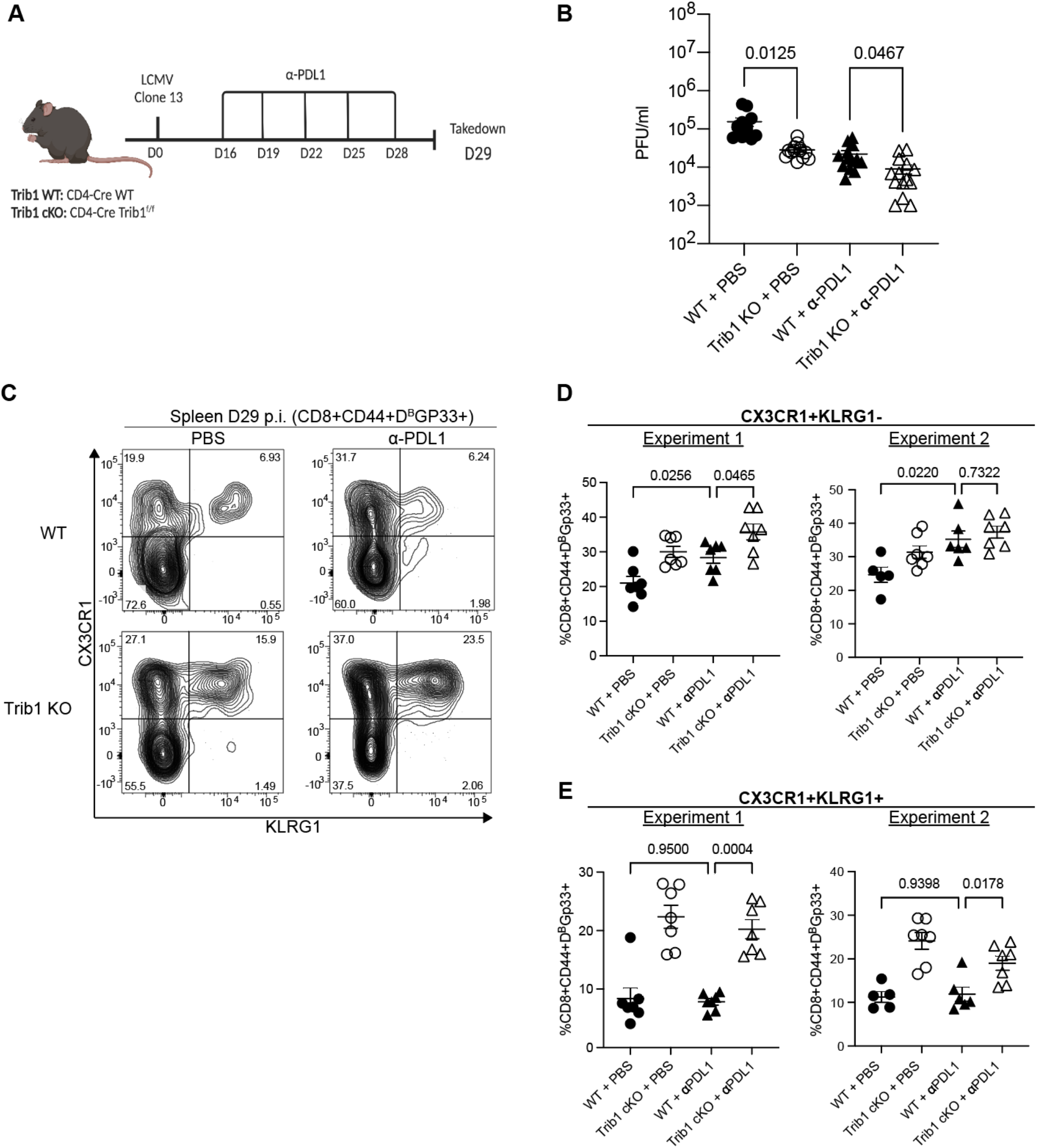
Trib1 deletion augments PDL1 blockade to reduce viral titers in chronic LCMV infection. (A) Experimental design. WT and Trib1 KO mice were infected with clone 13 and then treated with 5 doses of either PBS (control) or anti-PDL1 antibody every 3 days, beginning at day 16. Mice were taken down at day 29. (B) Viral titers from serum of Trib1 WT or KO mice treated with either PBS or anti-PDL1 as determined by plaque assay. (C) Representative flow plots with CD8^+^CD44^+^Gp33-Tetramer^+^ cells that are either CX3CR1^+^KLRG1^-^ T_INT_ or CX3CR1^+^KLRG1^+^ T_KLR_ at day 29 in the spleen. (D) Proportion of T_INT_ or (E) T_KLR_ as gated in (C). Data in (C-E) are representative of 2 experiments with 5-7 mice in each treatment group. Error bars are mean ± SEM. Data analyzed using a one-way ANOVA with Welch’s correction based on variance between samples and were adjusted for multiple comparisons. WT: CD4-cre^+^ Trib1^+/+^, Trib1 cKO: CD4-cre^+^ Trib1^F/F^

To investigate on a cellular level how Trib1 deletion impacts CD8^+^ T_EX_ during PD1 inhibition, CD8^+^CD44^+^Gp33^+^ splenocytes were analyzed for the presence of CX3CR1^+^ T_INT_ and KLRG1^+^ T_KLR_ subsets (**Fig. 5C**). As expected, anti-PDL1 treatment increased the frequency of CX3CR1^+^KLRG1^-^ T_INT_ cells in WT mice (**Fig. 5D**). Trib1 deletion combined with anti-PDL1 treatment also increased the T_INT_ compared to WT treated mice, but with varying significance (**Fig. 5D)**. Notably, anti-PDL1 treatment did not impact CX3CR1^+^KLRG1^+^ T_KLR_ in WT mice (**Fig. 5E**), and addition of anti-PDL1 treatment to Trib1 KO mice did not further increase the T_KLR_ population. Together, these data suggest that inhibiting Trib1 augments PD1 blockade by enhancing the anti-PDL1 expanded T_INT_ population while simultaneously supporting differentiation to the more effector-like T_KLR_ fate.

## Discussion

Here we demonstrate that Trib1 deletion from T cells during chronic infection increases the expression of KLR-associated genes at all stages of exhaustion and drives the clonal expansion and differentiation of T_INT_ towards a T_KLR_ identity. Little is known about the signaling events and molecules involved in terminal differentiation of exhausted CD8 T cells and even less is known about the signaling processes involved in pushing exhausted CD8 T cells towards a fate with enhanced effector function. We show that deleting a single signaling molecule, Trib1, promoted T_KLR_ differentiation to divert the pathway of exhaustion towards a more effector-like state, and that this skewing was accompanied by greater viral control. Furthermore, Trib1 deletion improved anti-PDL1 treatment in chronically infected mice, an outcome associated with expansion of both T_INT_ and T_KLR_, suggesting that a viable strategy for reinvigorating exhausted CD8 T cells is to shift the pathway of exhaustion towards a T_KLR_ fate. This finding is in accordance with recent reports that KLR-expressing CD8^+^ T cells are clinically important in several settings where exhaustion poses a significant barrier to T cell function ^17-20^, and that T_KLR_ may serve as an alternative CD8^+^ T cell fate to the canonical terminally exhausted T_TERM_ cell ^4,14,15^. This is especially relevant given that Trib1 is highly expressed in circulating human CD8^+^PD1^+^ exhausted T cells and has multiple ACRs in exhausted TILs isolated from human basal cell carcinoma ^25^. Our data in the setting of a well-established preclinical mouse model for T cell exhaustion in conjunction with the finding that Trib1 is highly expressed in human T_EX_ suggests that manipulating Trib1, or it’s signaling partners, could provide a novel means to improve CD8 T cell function in the setting of chronic infection or cancer therapy.

CD8 T cell expression of KLRs, canonical NK cell inhibitory and activating receptors, is well documented in acute infection, and reports on the role of KLRs during chronic infection and cancer, have been increasingly described ^4,14,17-20^. Recently, LCMV-specific KLR-expressing CD8^+^ T cells were identified during chronic LCMV infection ^4,14,15^. These cells were both transcriptionally and epigenetically distinct from other CD8 T cells present in either acute infection or earlier in chronic infection ^4,14^, indicating that they may share some overlapping biologic features with short-lived effector CD8 T cells but that their presence is distinct to the exhaustion milieu. Furthermore, these studies suggest that T_KLR_ differentiation is an alternative fate to terminally exhausted T_TERM_ cells downstream of CX3CR1^+^ T_INT_ cells. Our scRNA-sequencing and RNA velocity analyses confirm this bifurcation model and show that Trib1 is an important signaling protein that regulates the balance between T_KLR_ verses T_TERM_ differentiation. The mechanism by which Trib1 regulates this balance is at least two-fold. First, loss of Trib1 expands both the T_INT_ and T_KLR_ subsets with a concomitant decrease in T_TERM_ cells, suggesting that Trib1 deletion regulates the bipotential fate choice of T_INT_ by pushing them towards a T_KLR_ fate. This enrichment was reflected by cell numbers, especially within the T_KLR_ subset, and strongly evidenced by the clonal behaviors of individual TCR clones found in Trib1KO mice. In the absence of Trib1, clonally expanded CD8 T cells were heavily biased towards the T_INT_ and T_KLR_ subsets, indicating robust expansion and/or increased survival of expanded clones within these subsets. Second, Trib1 impacts the differentiation from the T_INT_ to more terminal subsets in a CD8 T cell-intrinsic manner. In co-adoptive transfer experiments of WT and Trib1KO P14 cells, the loss of Trib1 in CD8^+^ P14 cells resulted in a specific increase in T_KLR_ differentiation, indicating skewing to this subset was CD8 T cell-intrinsic. This is an important distinction, as CD4^+^ T cell help is known to enhance viral control and maintenance of the T_INT_ population ^13^, and Trib1 KO CD4+ T cells have increased cytokine production in this setting ^16^. Together, these features of Trib1 loss result in an exhausted CD8 T cell pool that is enriched for more effector-like cells. Though our studies were performed in the context of Trib1 deletion, our data suggest that Trib1 may actively promote terminal exhaustion and hinder viral control by inhibiting T_KLR_ differentiation and tipping the T_EX_ pathway towards the T_TERM_ fate.

Our data also show that deleting one signaling molecule enhances immune checkpoint blockade by shifting the exhausted CD8 T cell milieu towards a T_KLR_ fate. PD1 is a well-described inhibitory receptor that is upregulated beginning at the T_PROG_ stage, with highest expression on T_TERM_ and lowest expression on T_KLR_. Blockade of PD1 or its ligand PDL1 results in improved viral control during clone 13 infection, and in improved anti-tumoral immunity in both mouse and human cancers ^3,7-9,30^. Here, we demonstrate that Trib1 KO improves PD1 blockade by decreasing viral load compared to WT mice also treated with a blocking PD-L1 antibody alone, highlighting a potential therapeutic target for improving immune therapies for chronic infection or cancer. Additionally, while targeting either PD1 or Trib1 alone improves viral control, we demonstrate that Trib1 KO produced distinct effects on exhausted CD8 T cell ontogeny compared to anti-PD-L1 treatment alone. Specifically, PD1 blockade acts on the T_PROG_ subset to promote accumulation of CX3CR1^+^ T_INT 7-11_. It is important to note that these studies included all CX3CR1^+^CD8^+^ cells within the broad definition of intermediate or transitional cells and did not investigate the T_KLR_ subset ^10,12,13^. We also find that PD1 blockade leads to an expansion of the T_INT_ cells. Importantly, it is Trib1 deficiency and not PD1 blockade that augments the CX3CR1^+^KLRG1^+^ T_KLR_ population. Combining PD1 blockade and Trib1 deletion thereby enhances both T_INT_ and T_KLR_ subsets, producing a more robust restoration of viral control and further demonstrating the translational implications of targeting regulators of T_KLR_ differentiation to improve immune therapy.

In summary, we utilized multiple techniques to demonstrate that deleting Trib1 results in improved viral control that is specifically associated with T_INT_ enrichment and T_KLR_ expansion. Furthermore, Trib1 deletion can be utilized to further augment PD1 blockade in a model of chronic disease. Combined, these findings identify that Trib1 controls T_EX_ cell differentiation trajectories and suggest that Trib1 or other regulators of T_KLR_ differentiation may offer new strategies for improving immunotherapies such as ICB during chronic infection and cancer.

## Star Methods

### Mice

*Cd4-Cre*^+^*Trib1*^*f/f*^ Trib1 KO mice were generated as previously described ^16^, and were used with age and sex matched *Cd4-Cre*^+^ *Trib1*^+/+^ WT mice as controls. Both female and male mice were used in experiments. For adoptive transfer experiments, *Cd4-Cre*^*+*^ P14 mice with the WT Trib1 gene and a transgenic TCR specific for LCMV peptide gp33-41 were crossed with *Cd4-Cre*^+^*Trib1*^*f/f*^ Trib1 KO mice to generate conditional Trib1 KO P14 mice. All Trib1 WT and KO mice were bred and backcrossed on the C57BL/6 background. Recipient C57BL/6 CD45 congenic mice were purchased from Jackson Laboratory. Mice were housed in a pathogen-controlled environment and all breeding and animal use was performed in accordance with Institutional Animal Care and Use Committee (IACUC) guidelines at the University of Pennsylvania.

### LCMV Infection

For all experiments, 5–7-week-old mice were infected with LCMV clone 13 (4 × 10^6^ PFU) by tail vein injection. LCMV virus was grown in BHK cells and tittered by plaque assay in the laboratory of John Wherry at the University of Pennsylvania. For adoptive transfer experiments, recipient mice were infected one day after adoptive transfer of P14 cells as previously described.

### Adoptive T cell transfer

Donor PBMCs were isolated from WT or Trib1 KO P14 mice using Histopaque-1083 centrifugation (Sigma-Aldrich). A small aliquot of PBMCs was stained and analyzed by flow cytometry to determine the frequency of P14 T cells from each donor sample. Recipient mice were intravenously injected with donor PBMCs containing 500 WT and 500 Trib1 KO P14 cells. Mice were infected with LCMV clone 13 one day after adoptive cell transfer. Donor WT and Trib1 KO P14 mice and recipient mice were all congenically distinct based on the expression of CD45.1 and CD45.2.

### PDL1 Antibody Treatment

WT or Trib1 KO mice were infected with clone 13 on day 0. On day 15, all mice were bled and PD1 expression was confirmed on peripheral CD8^+^ T cells. Mice were given 5 doses of either 200ug anti-PDL1 antibody (10F.9G2, BioXCell) or sterile PBS by intraperitoneal injection every 3 days beginning at day 16. Mice were analyzed on day 29, one day after final treatment.

### Flow Cytometry and Cell Sorting

Spleens were manually disrupted by passing through a 70 uM cell strainer to generate a single cell suspension. Cells were resuspended in PBS then stained with the amine-reactive dye Live/Dead Aqua (Invitrogen) for 20 minutes. Surface staining was performed with antibody cocktails using antibodies for Ly108 (13G3, BD), CX3CR1 (SA11F11, Biolegend), CD4 (RM4-5 BD and Biolegend), CD44 (IM7, Biolegend, BD, or Invitrogen), CD8 (53-6.7, BD), KLRG1 (2F1, Biolegend), CD101 (Moushi101, eBioscience), TCRbeta (H57-597, Biolegend), NKG2A (16A11, Biolegend), CD94 (18d3, Biolegend), CD45.2 (104, Biolegend), and CD45.1 (A20, Invitrogen). Tetramer specific for H2-D^b^-restricted to the gp33-41 epitope of LCMV was obtained from the National Institutes of Health Tetramer Core Facility. Surface staining was completed in staining buffer of 1:1 Brilliant Stain (BD) and 2% FBS in PBS with 5% Fc block (2.4G) and stained for 1 hour at 4 degrees. Cells were washed and fixed in 4% paraformaldehyde for flow cytometric analysis on a BD Fortessa Flow Cytometer. Data were analyzed using FlowJo software (TreeStar, version 10.8.0). For all analysis, doublets were excluded. Cells were gated on live events using Live/Dead Aqua (Invitrogen). For scRNA-sequencing and scTCR-sequencing, cells were sorted on a BD FACSAria II cell sorter as previously described ^16^.

### Serum Plaque Assays for LCMV Viral Load

Serum was obtained from peripheral blood of infected mice. Ten-fold dilutions of serum were incubated on adherent Vero cells for 1h at 37^°^C. Vero cells were then overlaid with a 1:1 mixture of medium and 1% agarose. Cells were incubated at 37^°^C for 4d. Cells were then overlaid with a 1:1:20 mixture of medium:1% agarose:neutral red dye overnight. PFUs were then counted, and viral load calculated.

### scRNA-sequencing and scTCR-sequencing

scRNA-sequencing was performed on sorted activated T cells from WT and Trib1 KO mice infected with LCMV clone 13 for 15d as described ^16^ using the Chromium Single Cell 5′ Library & Gel Bead Kit (10x Genomics). scTCR-sequencing and library generation were performed simultaneously using the Chromium Single Cell V(D)J Enrichment Kit for Mouse T cells (10x Genomics). Libraries were sequenced on a NextSeq 550 using paired-end sequencing on one high output FlowCell (Illumina).

### Single-cell RNA-seq data analysis

#### Analysis using TooManyCells

Previously published scRNA-seq data of Trib1 control and cKO CD44+ TCRB+ T cells from d15 p.i with LCMV cl13 (GSE143802) ^16^ was processed using the methodology of Rome et. al and the scRNA-seq analysis program, TooManyCells (v2.2.0.0) ^26^, with default settings. Before analyzing the data, Scrublet (v0.2-1-0) ^31^ was employed to detect the presence of doublets. Overall, only four doublets were detected in total from Trib1 control and KO cellranger outputs. The TooManyCells “--draw-leaf” flag was used to overlay upper-quartile scaled gene expression across all trees. To generate a new tree containing only cells with low CD4 expression, CD4 expression was first projected across the tree with an exact gene expression cutoff of 0, defining CD4 “high” and “low” expression. A cell whitelist file containing barcodes of only CD4 low-expressing cells was then created with the TooManyCells “--labels-output” flag and used to generate a new tree with the TooManyCells flag “--cell-whitelist-file.” The tree was trimmed using the flag “--smart-cutoff 1,” which prunes nodes one standard deviation away from the median of all node sizes of the original tree. Differential gene expression between clusters was obtained using edgeR ^32^ through TooManyCells differential with the flag “--normalization ‘NoneNorm’.” Volcano plots and bar graphs were plotted using GraphPad Prism (v9.3.1).

#### GSEA Analysis

GSEA was performed using the Broad Institute software (https://www.gsea-msigdb.org/gsea/index.jsp). Previously published gene signatures ^4^ (consisting of 500 genes) were used as gene sets to compare to a log normalized count matrix with CD4 low only cells, annotated by TooManyCells cluster using a phenotype file. Genes in the count matrix were ranked by Signal2Noise and GSEA was performed using 1000 phenotype permutations.

#### Analysis based in Seurat

Cellranger outputs were additionally analyzed in R (v4.1.2) using Seurat (v4.1.0) ^33,34^. Cells with numbers of features over 2500 and below 200 and mitochondrial reads above 5% were removed. The remaining cells were assigned identities based on TooManyCells cluster barcodes and cell barcodes not found in the Cd4 low tree were subsetted out of the Seurat object. Gene expression was then normalized using Seurat’s NormalizeData function and scaled by the ScaleData function. The DoHeatmap function was used to generate a heatmap of the top 15 significant DEGs for each cluster, with average expression of genes per cluster generated by the AverageExpression function. Cluster projections of cluster 3 and cluster 9 was done following methodology from Giles et. Al ^4^ with S and G2M scores calculated for each cell using the CellCycleScoring function.

#### RNA Velocity Analysis

Velocyto (v.17.17) ^28^ was run on the cellranger counts output folder to generate a loom file containing the spliced/unspliced counts matrix. The loom file was merged to an annadata object containing metadata of either Trib1 WT or KO counts matrix restricted to barcodes found in the cell whitelist file using Python (v3.6.8) package scanPY (v1.9.1) ^35^. The annadata object was preprocessed in scanPY following Seurat convention. UMAPs of Trib1 WT and KO were created by seeing the number of neighbors to 100 and the numbers of PCs to 50. The spliced/unspliced counts were then normalized using the package scVelo (v0.2.5) ^27^ and genes without a shared splice count of 20 were filtered out of the annadata object. The moments were then computed across 30 neighbors and 30 PCs. Velocities were estimated using scVelo’s dynamical model, and velocities were then calculated and embedded onto UMAPs. The python package cellrank (v1.5.1) ^29^ was then used to calculate terminal states, initial states, macrostate lineages and site directed PAGA, with a probability threshold of .95.

### Single-cell TCR-seq data analysis

scTCR reads were aligned to the mm10 genome and consensus TCR annotations were performed using the cellranger vdj pipeline (10X genomics, v3.1.0) for Trib1 WT and cKO fastqs separately. TCR clonotype data and gene expression data of CD4 low expressing cells were combined into one overall Seurat object based on shared cell barcodes. Clonotypes with cell populations higher than 5 were used for downstream analysis. Stacked bar graphs of cluster counts per clonotype were visualized using dittBarPlot (dittoSeq, v1.6.0)^36^ and upset plots were created using UpsetR (v1.4.0) ^37^

## Data and Code Availability

All scRNA-seq and scTCR-seq data generated in this study will be deposited on GEO prior to publication. No new algorithms were developed during this study. Analyses were performed as described and scripts will be made available upon request.

## Statistics

Statistical methods for each experiment are briefly described in the corresponding figure legend. For direct comparisons of flow cytometry data or viral load, comparisons of two groups are calculated using a student t-test. If variance is statistically significant, a t-test with Welch’s correction was used. For P14 adoptive transfer experiments, a paired t-test was used given that the comparisons were calculated based on two populations within the same recipient mouse. For comparisons involving more than two groups, a one-way ANOVA was used with Welch’s correction based on variance between samples and were adjusted for multiple comparisons. Statistical methods for computational analysis is described in the corresponding methods section.

## Study Approval

No human data was included in this manuscript. Animal studies were performed with the approval of the University of Pennsylvania Institutional Animal Care and Use Committee (IACUC) for the described experiments.

## Acknowledgments

All flow cytometry in this manuscript was performed using the University of Pennsylvania Flow Cytometry and Cell Sorting Core (National Institutes of Health P30CA016520), and mice were bred and housed in the University of Pennsylvania mouse husbandry core. Additionally, we thank, GW Schwartz (University of Pennsylvania, Philadelphia, PA) and Ansuman Satpathy, Bence Daniel, and Joy Pai (Stanford University, Palo Alto, CA) for advice regarding scRNA-sequencing and scTCR-sequencing analyses. We also thank the Pear and Jordan labs for thoughtful discussions. Schematics of T cell exhaustion were created using BioRender.

This study was supported by grants from National Institutes of Health grants R01AI047833 (MSJ and WSP), NIH grants AI155577, AI115712, AI117950, AI108545, AI082630 and CA210944 (EJW), T32CA009615 (SEM), K12CA076931 (SEM), T32CA009140 (KSR and JRG), P30CA016520 (PAG), and a Cancer Research Institute-Mark Foundation Fellowship (JRG). Work in the Wherry lab is supported by the Parker Institute for Cancer Immunotherapy.

Declaration of interests: EJW is a member of the Parker Institute for Cancer Immunotherapy which supports work in the Wherry Lab. EJW is an advisor for Danger Bio, Marengo, Janssen, New Limit, Pluto Immunotherapeutics, Related Sciences, Rubius Therapeutics, Synthekine, and Surface Oncology. EJW is a founder of Surface Oncology, Danger Bio, and Arsenal Biosciences.

## Supplementary Materials

**Figure S1:**
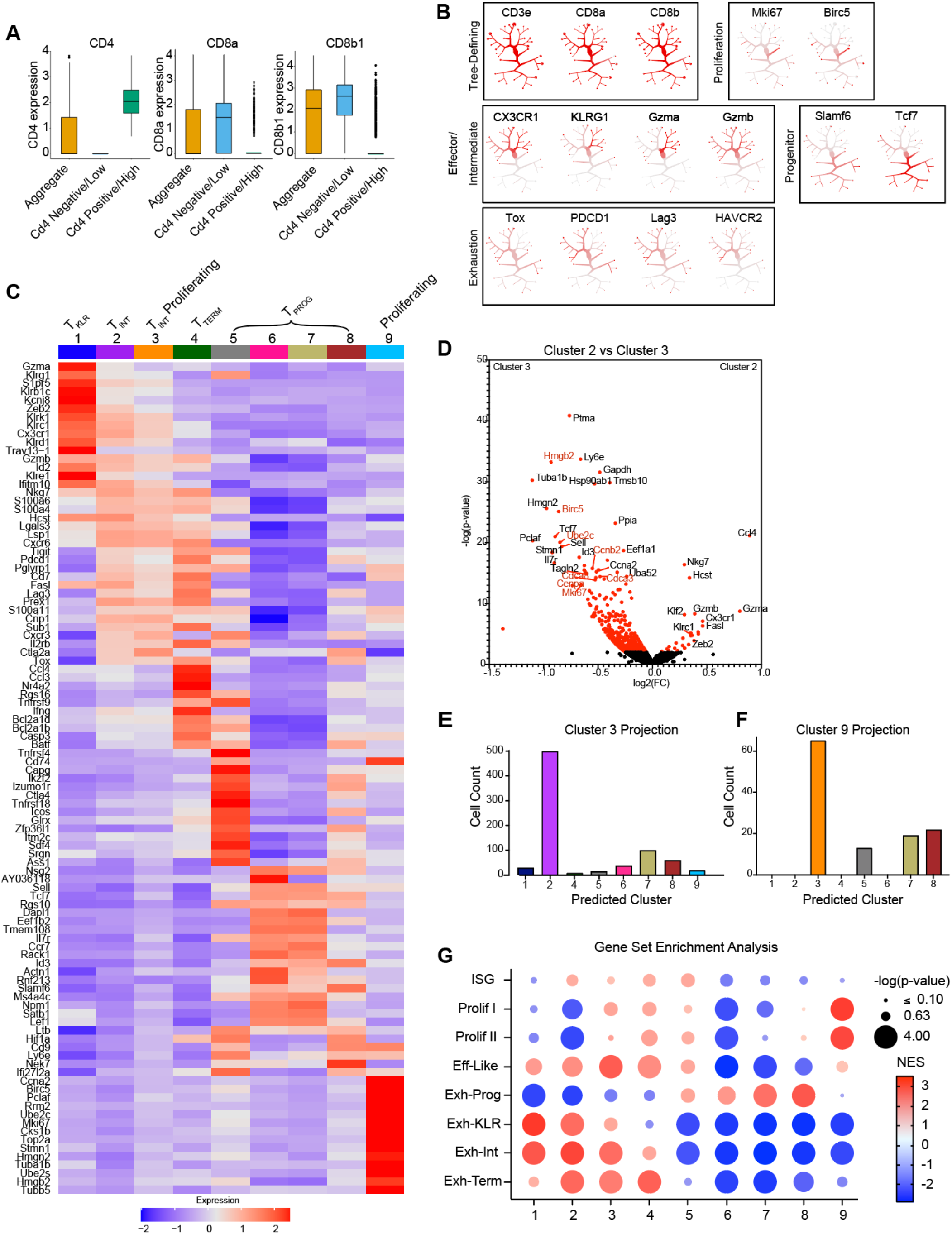
A complete pathway of CD8^+^ T_EX_ differentiation can be identified in both Trib1 WT and Trib1 KO mice. (A) Relative expression of *CD4, CD8a*, and *CD8b* mRNA in all aggregate cells sorted for scRNA-seq as referenced in Rome et al^16^ (yellow), in the *CD4*-low tree in Figure 1b (blue), or in the *CD4*-postive/high cells which were excluded from the tree in Figure 1b (green). (B) Normalized reference gene expression on the CD8^+^ tree, where red = higher expression and gray/white = lower expression. (C) Heatmap demonstrating top DEGs within each of 9 clusters, where red = higher expression within that cluster and blue = lower expression. (D) Volcano plot of differentially expressed genes between Cluster 2 and Cluster 3. Proliferation genes are highlighted in red font. (E) Predicted identity of cluster 3 cells when proliferating genes were regressed out of analysis using the SCTransform function in Seurat. (F) Predicted identify of cluster 9 cells when proliferating genes were regressed out of analysis using the SCTransform function in Seurat. (F) GSEA analysis comparing the 9 clusters identified in this paper (x-axis) to the 8 populations identified at day 15 of LCMV clone 13 infection by Giles et al ^4^ (y axis). Trib1 WT: CD4-cre^+^ Trib1^+/+^, Trib1 KO: CD4-cre^+^ Trib1^F/F^

**Figure S2:**
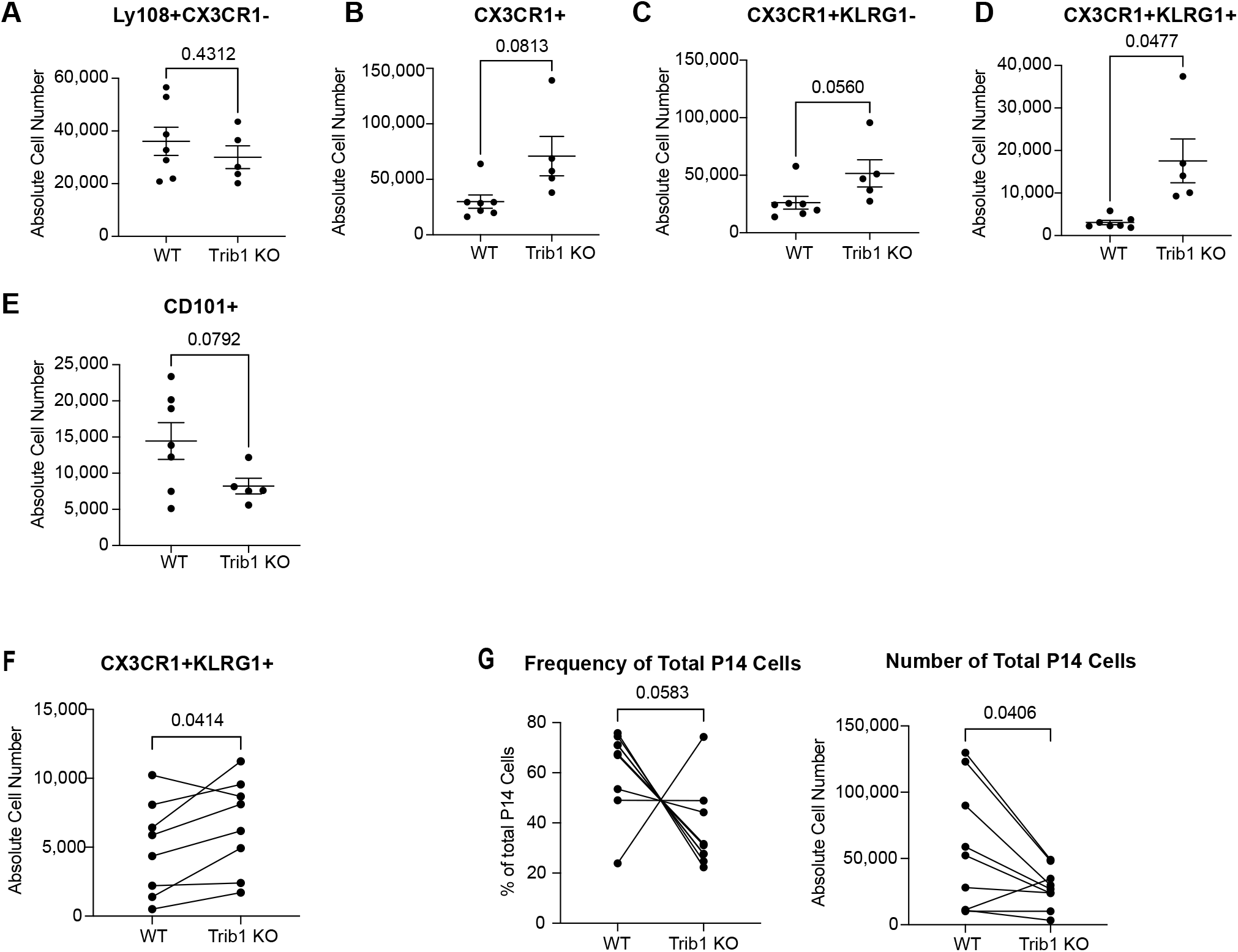
Trib1 restrains the absolute number of T_KLR_ that persist during chronic infection. (A-E) Absolute numbers of the indicated population from the same experiment shown in Figure 2C-E. For this analysis, cells were gated on CD8^+^CD44^+^D^B^Gp33^+^ events as shown in Figures 2C-E. (F) Absolute number of CX3CR1^+^KLRG1^+^ P14 cells in the adoptive transfer experiments described in Figure 2G-H. Events are gated on CD8^+^D^B^Gp33^+^ cells and the appropriate congenic marker for WT (CD4.1/2) or KO (CD45.1) cells to identify genotype-specific P14 cells. (G) Frequency of total P14 cells by genotype as a percentage of total P14 cells or as absolute number. For A-G, absolute cell numbers were calculated by multiplying the frequency of each gated population as a percentage of live cells and by total live splenocytes for each mouse. For A-E, Error bars are mean ± SEM. P values calculated using either Student’s t test or unpaired t test with Welch’s correction based on variance between genotypes. For F-G, P values were calculated using a paired T test given that WT and KO cells were co-transferred into the same individual recipients. Trib1 WT: CD4-cre^+^ Trib1^+/+^, Trib1 KO: CD4-cre^+^ Trib1^F/F^

